# Mitochondria-Associated Membranes Are Not Altered In Immune Cells In T2D

**DOI:** 10.1101/2024.03.25.586170

**Authors:** Samantha N. Hart, Raji Lenin, Jamie Sturgill, Philip A. Kern, Barbara Nikolajczyk

## Abstract

Metabolism research is increasingly recognizing the contributions of organelle crosstalk to metabolic regulation. Mitochondria-associated membranes (MAMs), which are structures connecting the mitochondria and endoplasmic reticulum (ER), are critical in a myriad of cellular functions linked to cellular metabolism. MAMs control calcium signaling, mitochondrial transport, redox balance, protein folding/degradation, and in some studies, metabolic health. The possibility that MAMs drive changes in cellular function in individuals with Type 2 Diabetes (T2D) is controversial. Although disruptions in MAMs that change the distance between the mitochondria and ER, MAM protein composition, or disrupt downstream signaling, can perpetuate inflammation, one key trait of T2D. However, the full scope of this structure’s role in immune cell health and thus T2D-associated inflammation remains unknown. We show that human immune cell MAM proteins and their associated functions are not altered by T2D and thus unlikely to contribute to metaflammation.

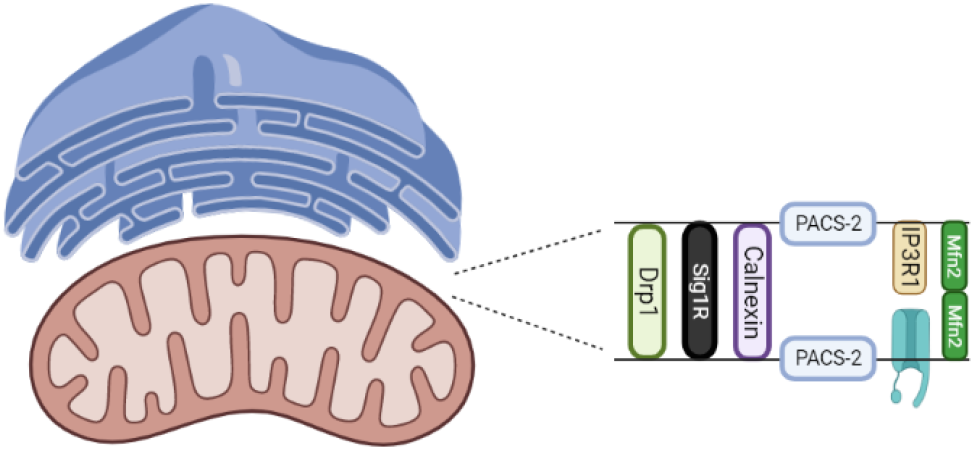

## INTRODUCTION

Mitochondria play important roles in inflammation and obesity/Type 2 Diabetes (T2D) due to their role in energy production, proliferation, and metabolite generation [25]. Disruptions in mitochondrial function increases oxidative stress [24], and in the case of T2D, mitochondrial dysfunction in immune cells can induce IL-1β release, decreasing sensitivity to insulin [38]. Mitochondrial dysregulation in adipose-associated immune cells and adipocytes increases superoxide production and decreases ATP production due to inefficient nutrient oxidation [19], fueling a feed-forward loop of decline in mitochondrial health in those with obesity and T2D.

The endoplasmic reticulum (ER) is similarly central to cellular health; it is responsible for the storage and export of calcium ions, production of phospholipids, and protein production/folding [12, 30]. The ER is tethered to the mitochondria by a complex, multi-protein scaffold termed mitochondria-associated membranes (MAMs) that regulates the distance between the mitochondria and ER to control inter-organelle communication and the related transport of ions/proteins. MAMs are comprised of over 1,000 proteins [39], some of which have been identified as important in cell signaling, transport, intracellular communication, and distance between organelles. Malfunction of MAM protein(s) compromises communication across these structures [9], given that MAMs provide a network for transportation of key molecules between the mitochondria and ER [14,15, 28]. Calcium (Ca^2+^), a foundational signaling molecule, enters the mitochondria from the ER mainly through MAMs. IP3R1 is a central protein within MAMs that directly impacts transport of Ca^2+^ ions by sensing cytosolic concentrations and releasing appropriate amounts of Ca^2+^ out of the ER. In turn, Ca^2+^ is brought into the mitochondria through voltage-dependent anion channels (VDACs) [10].

In addition to IP3R1, MAMs include PACS-2, Drp1, Sigma1, and Mfn2 [21]. These proteins collaborate to transport Ca^2+^ and other small molecules from the ER to the mitochondria [23]. For example, Sigma1 binds to IP3R1 to facilitate Ca^2+^ transport [6, 11]. Drp1 is important not only in MAMs but also for mitochondrial fission and apoptosis. This multifunctionality provides a challenge for elucidating Drp1’s contribution to MAM function [2].

MAMs are involved in a myriad of inflammatory processes including IL-2 production/secretion, which has been linked to Ca^2+^ flux [23]. MAM-associated proteins contribute to glucose homeostasis, and thus MAM dysfunction may contribute to the development of T2D [31]. MAMs are also involved in insulin signaling, in part because some proteins that connect the ER to the mitochondria are also signaling proteins for insulin production i.e., PACS-2 and HK2 [34]. In immune cells, MAMs are important for signaling involved in initiating the immune response [32]. The literature on MAMs’ contribution to obesity and insulin resistance generally (though not always) indicates that compromised MAMs negatively impact metabolic health. These include studies showing that MAM alterations in ß cells associate with the development of insulin resistance [40], and studies showing that MAM disruption in skeletal muscle precedes developing insulin resistance. Therefore, dysfunctional MAMs, at least in classical metabolic regulatory tissues, may be viable markers for likely T2D progression [35].

Cells/tissues from individuals with chronic hyperglycemia, a clinical indicator of metabolic disease, produce more reactive oxygen species (ROS) [7] as a byproduct of mitochondrial respiration. ROS is both a cause and effect of inflammation and multiple T2D comorbidities. Unmitigated ROS production leads to mitochondrial stress, DNA damage, and disturbances in signal transduction [18], all of which can fuel chronic inflammation when anti-oxidants are insufficient either in immune cells or the broader T2D milieu [3, 22, 29]. Oxidative stress, a common outcome of higher ROS, is often increased in cells with MAM alterations, which in turn contributes to chronic comorbidities of T2D [3, 36]. Obesity/T2D-associated oxidative stress fuels a feed-forward cycle of ER and mitochondria stress through MAMs to perpetuate stress-mediated dysfunction in many cell types and tissues (Vance et al, 2014), although direct links to T2D-associated inflammation remain untested.

Our new work shows that MAMs are not significantly different in immune cells from those with obesity and either T2D or normal glucose tolerance (NGT). MAM-associated protein abundance and total ROS production did not differ in immune cells from the two cohorts, although cells from T2D subjects had higher amounts of mitochondrial superoxide than the NGT cohort. Exposing cell lines to hyperglycemia similarly failed to change MAM-associated protein abundance. We conclude human immune cell MAMs are unlikely to contribute to metaflammation.

## METHODOLOGY

### Human subjects

This study had approval from Boston University and the University of Kentucky Institutional Review Boards in accordance with the Declaration of Helsinki.

One subject with T2D (as defined by American Diabetes criteria) was recruited from the Endocrinology clinic and the Center for Endocrinology, Diabetes and Nutrition at the Boston University Medical Center (BUSM) between January 10^th^, 2017 and January 10^th^, 2018. Subjects were also recruited from the central Kentucky area by the University of Kentucky Center for Clinical and Translational Sciences between September 1^st^, 2021 and January 26^th^, 2023. No recruiting took place between January 2018 and September 2021. The characteristics of the cohort used in this study are summarized in table 1. Total *n*=16; of those, 8 subjects had NGT and 8 subjects had T2D. Blood was drawn after >12 hours of fasting, and total PBMCs were isolated as we published (Nicholas et al, 2019). 56% of our cohort were male and 44% were female.

**Table 1:** Demographics of the cohort used for PBMC experiments. T2D status was clinically diagnosed. ж Indicates a significant difference between NGT and T2D subjects based on one-way ANOVA (p < 0.05).

|  | OBESE + NGT | OBESE + T2D |
| --- | --- | --- |
| AGE (YEARS) | 50.54 (41-60) | 47.46 (40-58) |
| BMI (KG * M<sup>2</sup>) | 35.23 (28-43.03) | 34.89 (29.5-38.06) |
| HBA1C % | 5.22 (4.9-5.5) | 9.01 (7.5-10.8) * |
| TOTAL N | 8 | 8 |
| FEMALES | 6 | 3 |
| MALES | 2 | 5 |
| AFRICAN DESCENT | 0 | 1 |
| WHITE/NON-HISPANIC | 8 | 6 |
| HISPANIC | 0 | 1 |

### PBMC Culture

500,000 PBMCs were cultured at 37°C (5% CO_2_) in RPMI 1640 medium with L-glutamine and 25mM HEPES (Gibco, Cat. No. 22400-089) under two conditions: unstimulated, or stimulated with anti-CD3/CD28 DynaBeads™ (Thermo Fisher, Cat. No. 11131D) at 1 bead/cell. All cells were cultured for 40 hours prior to analyses described below.

### Western Blots

Total protein was isolated from PBMC pellets using 30μL lysis buffer (Cell Signaling, Cat. No. 9803S) on ice for 30 minutes. Protein concentration was quantified by BCA assay. Twenty micrograms of protein was run on a 4-20% gradient precast gel (Bio-Rad, Cat. No. 456-1095) for separation, then transferred onto a polyvinylidene difluoride (PVDF) membrane for 100 minutes at 100 volts and 4°C. Primary antibodies against IP3R1, PACS-2, Mfn2, Drp1 and Sig1R were diluted to 1:500 in 3% BSA (Cell Signaling, Cat. No. A3294-100G) in TBST. β-Actin-specific antibody acted as a positive control diluted 1:1000 in 3% BSA.

### High Glucose Studies

Glucose-free RPMI media was supplemented with 5mM, 11mM, or 25mM glucose to represent normo- and hyper-glycemia. Although 25mM glucose is super-physiological, this condition was chosen to account for the higher glucose demands of cell lines for these experiments, in contrast to primary human cells used for other assays. Jurkat T cells (Cat. No. ATCC TIB-152) +/- CD3/CD28 DynaBeads and Raji B cells (Cat. No. ATCC CCL-86) +/- 250ng CpG (Cat. No. ODN 2216) were cultured for 40 hours before isolating proteins for analysis via Western Blot.

### Ca^2+^ Flux Measurement

250,000 PBMCs were cultured in 150μL media +/- CD3/CD28 DynaBeads for 40 hours. Cells were then collected and washed with DPBS (Gibco, Cat. No. 14190144), then resuspended in 250μL 1X DPBS. Fluo-4AM (Thermo Fisher, Cat. No. F14201) was added at a final concentration of 1nM, then the cells were incubated at 37°C (5%CO_2_) for 1 hour. Cells were pelleted then washed with 1X DPBS and resuspended in 250μL Hank’s balanced salt solution (Cat. No. J67681.K2). Labeled cells were loaded onto a black well, clear bottom 9-well plate and fluorescence was measured at 488nm (excitation) and 520nm (emission) in a micro plate reader.

### ROS Measurement

250,000 PBMCs were cultured in 150μL media +/- CD3/CD28 DynaBeads for 40 hours. Cells were collected, washed in 1X DPBS, then resuspended in 1X DPBS. Dyes for specific ROS species were then added in individual wells to the cells and incubated for 20-30 minutes at 37°C (5%CO_2_). Total superoxide was measured with Dihydroethidium (Thermo Fisher, Cat. No. D11347), mitochondrial superoxide was measured with mitoSOX (Thermo Fisher, Cat. No. M36008), and total peroxide was measured with CM-H2DCFDA (Abcam, Cat. No. ab113851).

## RESULTS

We used Western Blots to quantify MAM proteins in PBMCs from age- and BMI-matched subjects that differed based on a clinical T2D diagnosis (Table 1). While there are over 1,000 proteins within MAMs, we focused on IP3R1 and PACS-2, which have been demonstrated to contribute to MAM formation and function. IP3R1, PACS-2 and Sig1R expression was unchanged by stimulation, and did not differ in cells from the NGT compared to the T2D cohort (Fig. 1A & B). PBMCs from those with NGT had an upward trend in expression of MitoFusin2 (Mfn2) Dynamin-related protein 1 (Drp1) (p= 0.06 and p=0.08, respectively) when stimulated. Mfn2 and Drp1 increased in expression in cells from those with T2D when stimulated (p<0.001 for both proteins). However, there was no difference in Mfn2 or Drp1 expression between cohorts (Fig. 1B). These data support the conclusion that MAM protein expression is not impacted by T2D status.

**Figure 1:**
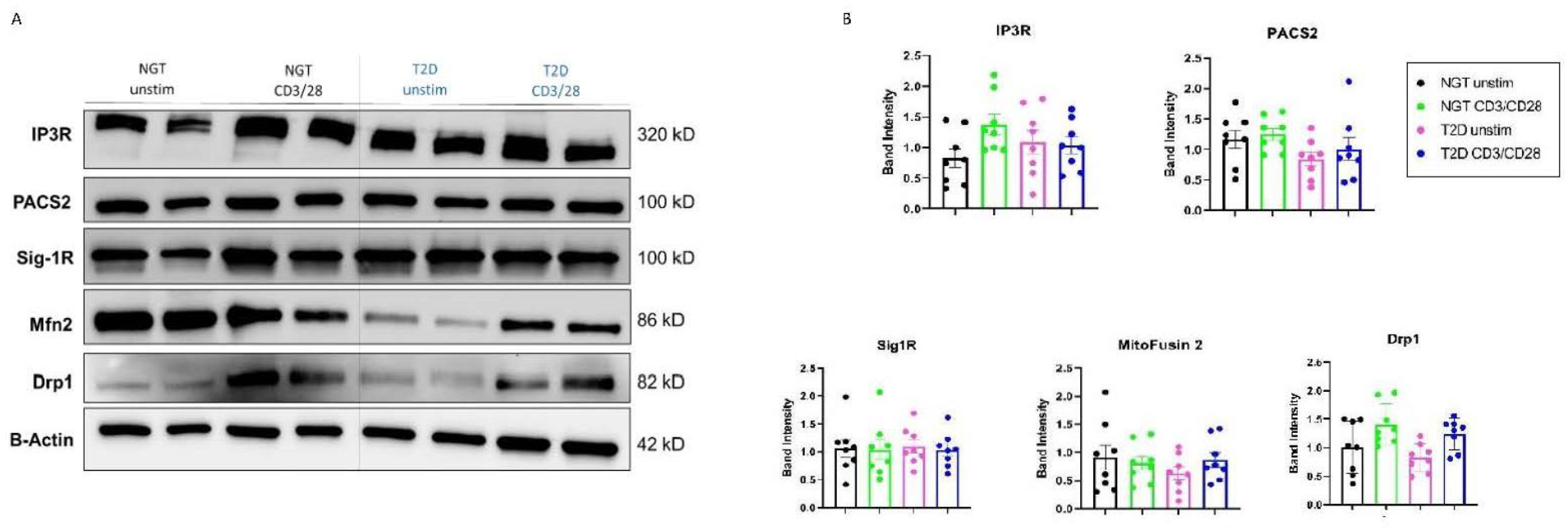
MAM proteins in PBMCs are not affected by stimulation or T2D status. A. Representative Western Blot, showing IP3R1, PACS2, Sig-1R, MitoFusin2, and Drp1 expression in PBMCs as indicated. *N=8* for NGT and T2D cohorts. B. Quantification of protein expression of blots (n = 8) represented in panel A. Differences between cohorts were determined via two-way ANOVA analysis. Error bars indicate SD.

To investigate the impact of hyperglycemia, a key characteristic of T2D, on MAM formation/function in immune cells, we assessed MAM protein expression in the Jurkat T cell line cultured under hyperglycemic conditions. 3-O-methyl D-glucose, a glucose analog, was used as a non-metabolizable osmolarity control. Hyperglycemia had a variable impact on MAM-associated proteins (Fig. 2). IP3R1 expression increases from normoglycemia to physiological hyperglycemia (5mM to 11mM glucose), then decreases in abundance with supraphysiological glucose (25mM). Sig1R expression is not significantly altered, although there is a trend towards lower Sig1R expression as glucose concentration increases. PACS2 and Mfn2 expression both have an upward trend between normoglycemia and physiological hyperglycemia, and supraphysiological hyperglycemia triggers significantly higher expression of these two proteins as compared to normoglycemia (Fig. 2B). Drp1 had similar expression when exposed to 5mM and 11mM glucose, but decreased significantly when exposed to 25mM glucose. A complete lack of glucose and/or the 3-O methyl D-glucose (osmolarity control) had an undefined impact on the cytoskeleton and likely overall cell health, indicated by the very low expression of ß-actin. Therefore, these data were not subjected to analysis. We conclude that hyperglycemia impacts MAM proteins non-uniformly.

**Figure 2:**
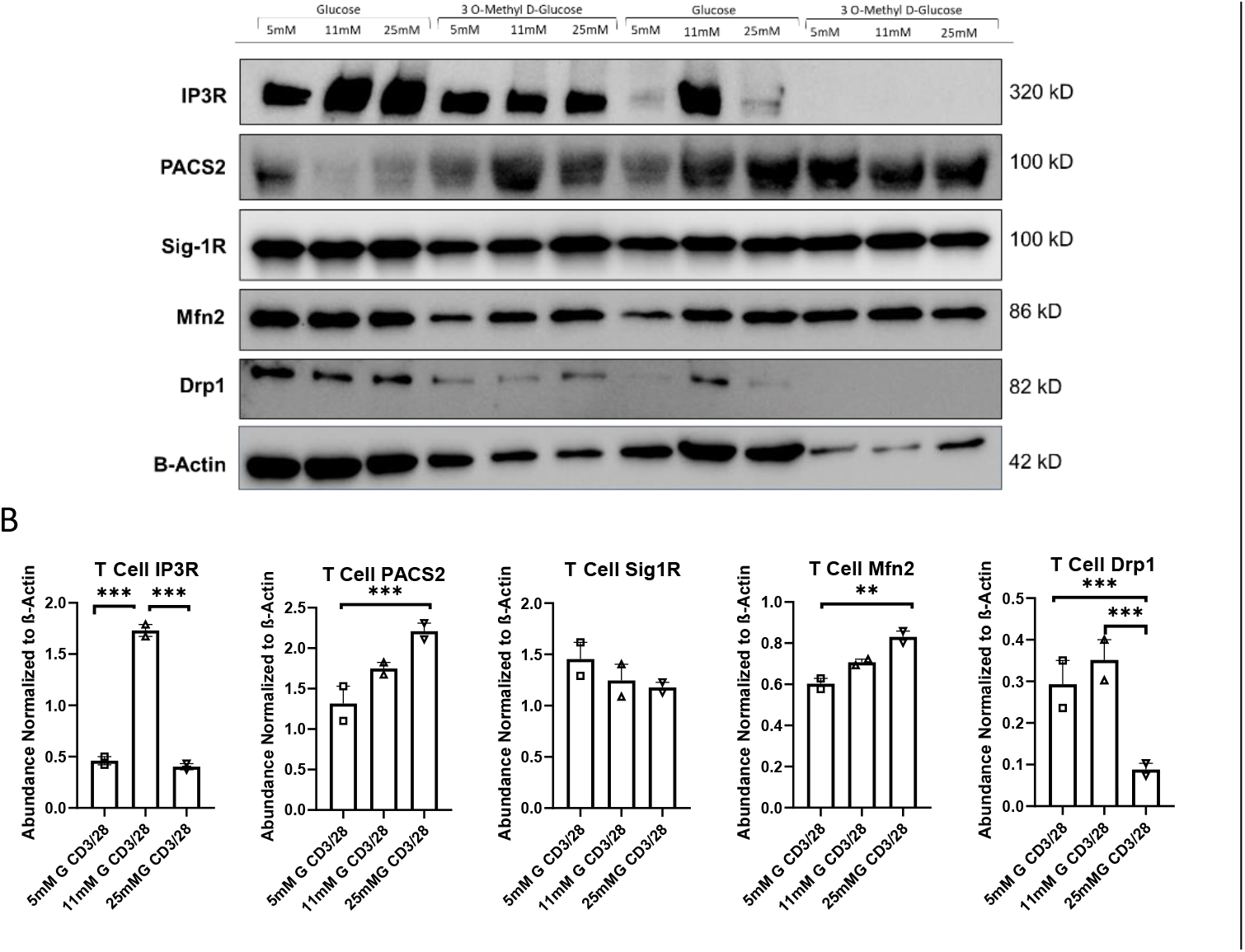
Response to high glucose in Jurkat T cells is non-uniform. A: Representative western blot of Jurkat T Cells in 5mM, 11mM, and 25mM glucose media. *N*=2 technical replicates B: Quantification of MAM-associated protein expression in Jurkat T cells. Differences in protein expression were determined via UNIANOVA analysis, and significance was accepted at p < 0.05 indicated by **. p < 0.01 indicated by ***. Error bars indicate range.

We similarly tested the impact of hyperglycemia on Raji B cells as a model of a second cell type that contributes to peripheral inflammation in T2D [17]. IP3R1, the master regulator of MAMs and primary importer of Ca^2+^ into the mitochondria, is unchanged by hypoglycemia. PACS2 expression by Raji B cells is similarly unchanged by hypo- or hyperglycemia (Fig. 3 A&B). In contrast to the Jurkat T cells, the osmolarity control (3 O-methyl D-glucose) modestly impacted ß-actin expression in B cells. Therefore, we could conclude that Sig1R expression was higher in 5mM, 11mM, and 25mM glucose compared to the glucose analog at the same concentrations.

**Figure 3:**
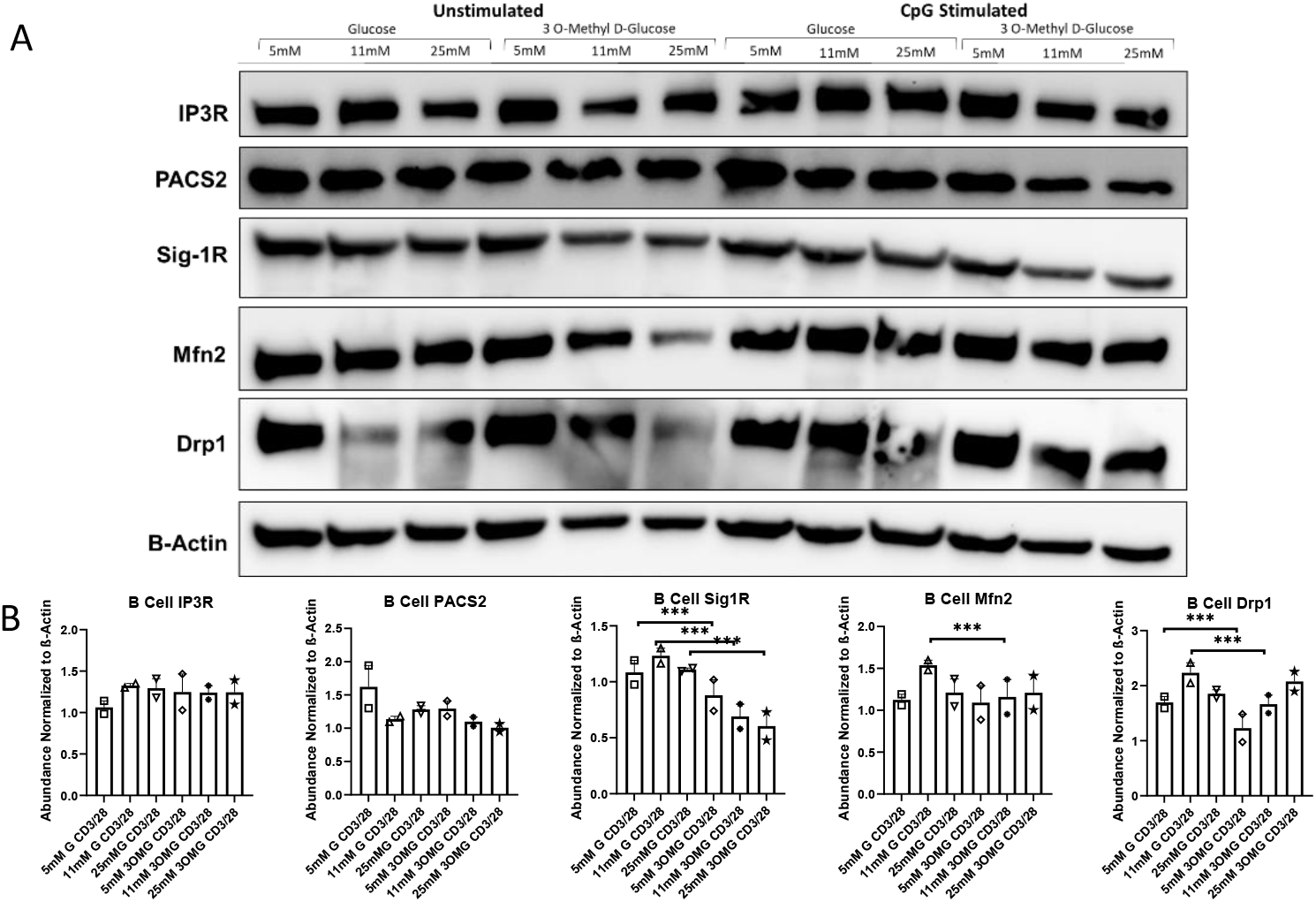
B Cell Line MAM expression is independent of stimulation and hyperglycemia. A: Representative Western Blot of Raji B cells +/- CpG (250ng/mL), grown glucose or osmolarity control as indicated. B: Quantification of A, with β-Actin as the control. Differences in protein expression were determined via UNIANOVA analysis, and significance was accepted at p < 0.05 (indicated by ***). Error bars indicate range.

**Figure 4:**
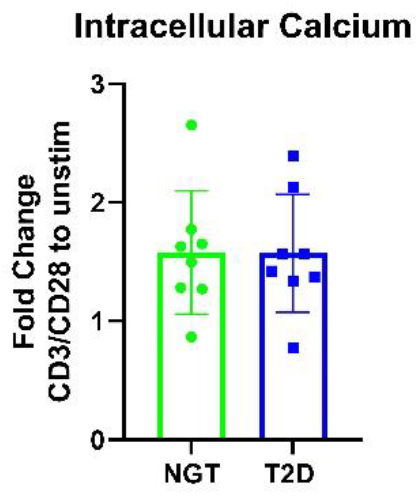
Intracellular Ca^2+^ flux is not different in NGT vs. T2D PBMCs. Plot of average +/- SD fold change between Fluo-4 AM fluorescent signal in stimulated versus unstimulated PBMCs from NGT and T2D subjects. P = 0.9369. Error bars indicate SD.

However, Sig1R expression was similar amongst glucose concentrations. Mfn2 and Drp1 were similarly higher in cultures with some glucose concentrations compared to the glucose analog controls, and has no significant differences in expression due to changes in media glucose. We conclude, as in T cells, hyperglycemia impacts MAM protein expression non-uniformly.

Because MAM protein abundance may not entirely reflect MAM function, we measured total intracellular Ca^2+^ concentrations as an indication of IP3R1 activity in primary human immune cells. PBMCs from those with T2D versus NGT had no difference in intracellular Ca^2+^. This functional indicator supported our conclusion that MAMs in immune cells do not change with T2D status.

ROS is typically higher in cells from people with T2D [7]. However, if MAMs are key regulators of ROS in immune cells, the similarities in MAM abundance/function in PBMCs from T2D versus NGT suggests that ROS may be similar in cells from these two cohorts. To investigate the importance of T2D on ROS accumulation in immune cells amidst similar MAM protein abundance, we quantified mitochondrial and cytosolic ROS concentrations in PBMCs. Total peroxide and total superoxide abundance in stimulated PBMCs were similar between cohorts (Fig. 5A and 5B). In contrast, mitochondrial superoxide was higher in stimulated PBMCs from T2D compared to NGT subjects (Fig. 5C). These data are consistent with previous demonstrations of generally elevated ROS in T2D, but suggests that differences in MAM abundance or function do not mechanistically explain such redox imbalance in immune cells.

**Figure 5:**
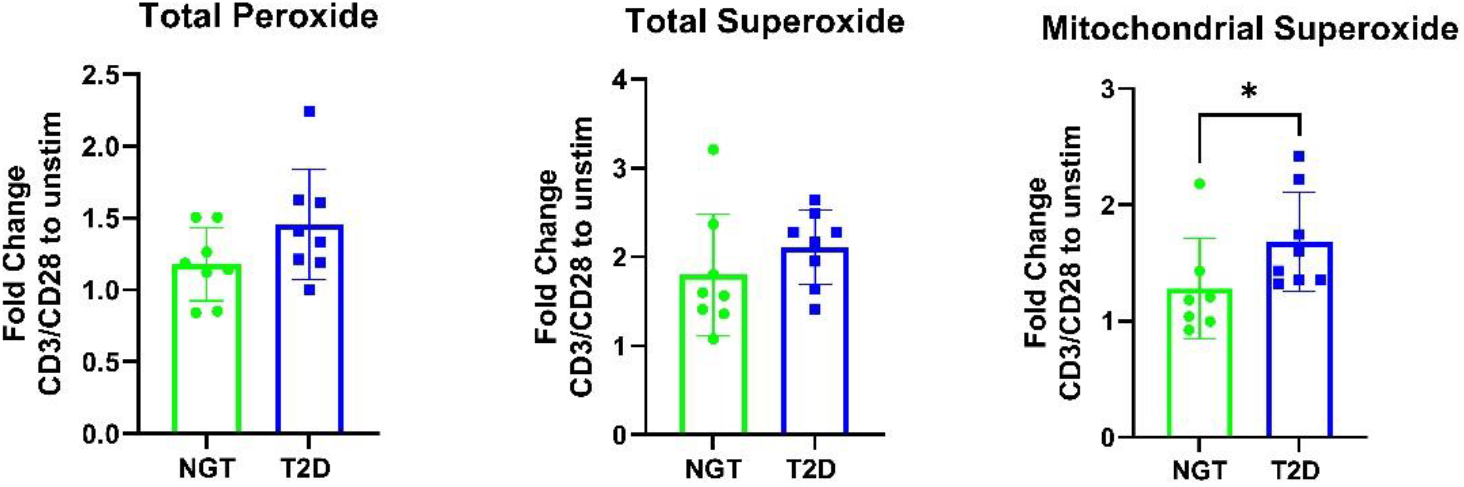
Mitochondrial superoxide is higher in PBMCs from T2D versus NGT subjects. A: Total peroxide, quantified by CM-H2DCFDA fluorescence. Error bars indicate SD. B: Total superoxide, quantified by dihydroethidium fluorescence. Error bars indicate SD. C: Mitochondrial superoxide, quantified by mitoSOX fluorescence. Error bars indicate SD.

## DISCUSSION

The full impact of T2D on immune cells and thus mechanistic explanations for inflammation-fueled comorbidities continues to be elucidated. Our new data showing that MAM protein abundance does not change in immune cells from individuals with T2D, and that the functions of those proteins are also not impacted by T2D status indicated that MAMs are highly unlikely to uniquely contribute to peripheral inflammation in T2D. This conclusion does not conflict with previous studies showing that MAMs are critical for extracellular signaling, particularly in the presence of inflammation [16, 20].

Our observation that IP3R1 expression and Ca^2+^ transport in immune cells was similar between cohorts independently supports the conclusion of T2D-independent expression and function of MAM proteins. We did not test the possibility that other Ca^2+^ transporters, like VDACs, compensate for theoretically possible changes in IP3R1 function not reflected by changes in protein expression and intracellular Ca^2+^ abundance.

Although T2D is characterized by hyperglycemia, hyperlipidemia, hyperinsulinemia, and a host of additional physiological changes, we focused on hyperglycemia as the clinically relevant indicator for T2D and mediator of T2D-associated inflammation [26, 37]. Our interpretation that hyperglycemia does not change MAM protein expression in immune cell lines is made cautiously, as we exposed the cells to the defined hyperglycemic conditions for days rather than years, as opposed to the long-term and erratic hyperglycemia characteristic of T2D, especially through sleep/wake cycles and circadian rhythm disturbances [13, 27].

Previous studies have shown that ROS production increases with cellular stress [33], usually as a byproduct of aberrant mitochondrial respiration [41], and its accumulation is detrimental to overall cellular health [1]. Although all of these mechanisms have been linked to oxidative stress in T2D [4, 5, 8, 22], changes in MAM structure/function are unlikely to contribute to T2D-associated ROS in immune cells. Because ROS activates inflammatory cytokine release from immune cells, we can also conclude this feed-forward loop explaining metaflammation is largely independent of MAM function. Instead, studies focused on changes in antioxidants or other redox-maintaining mechanisms, which we did not quantify in this study, may be more fruitful.

Overall, we have shown that abundance of key MAM proteins is not altered in the immune cells of those with T2D, correlating with no difference in intracellular calcium, one signature function of MAMs. We conclude that MAMs within immune cells are not different between NGT and T2D cohorts, and therefore not a likely contributor to the inflammatory phenotype typical of T2D status. Similar findings in T and B cell lines exposed acutely to hyperglycemia support our conclusion that MAM abundance is not significantly impacted by a key characteristic of T2D.

## ACKOWLEDGEMENTS

Funding for this project was provided by a pilot grant through the Diabetes Research center at Washington University, St. Louis, NIH P30 DK020579, a training grant through the University of Kentucky’s Center for Clinical and Translational Sciences, TL1001997, and the NIH National Center for Advancing Translational Sciences through grant number UL1TR001998. The content is solely the responsibility of the authors and does not necessarily represent the official views of the NIH.

## REFERENCES

1. Acker, T., et al. (2006). “The good, the bad and the ugly in oxygen-sensing: ROS, cytochromes and prolyl-hydroxylases.” Cardiovascular Research 71(2): 195–207.

2. Arasaki, K., et al. (2015). “A role for the ancient SNARE syntaxin 17 in regulating mitochondrial division.” Dev Cell 32(3): 304–317.

3. Auten, R. L. and J. M. Davis (2009). “Oxygen toxicity and reactive oxygen species: the devil is in the details.” Pediatr Res 66(2): 121–127.

4. Bartok, A., et al. (2019). “IP3 receptor isoforms differently regulate ER-mitochondrial contacts and local calcium transfer.” Nature Communications 10(1): 3726.

5. Boehning, D., et al. (2001). “Molecular Determinants of Ion Permeation and Selectivity in Inositol 1,4,5-Trisphosphate Receptor Ca2+ Channels*.” Journal of Biological Chemistry 276(17): 13509–13512.

6. Bonekamp, N. A., et al. (2021). “High levels of TFAM repress mammalian mitochondrial DNA transcription in vivo.” Life Sci Alliance 4(11).

7. Busik, J. V., et al. (2008). “Hyperglycemia-induced reactive oxygen species toxicity to endothelial cells is dependent on paracrine mediators.” Diabetes 57(7): 1952–1965.

8. Csordás, G., et al. (2012). “Calcium transport across the inner mitochondrial membrane: molecular mechanisms and pharmacology.” Mol Cell Endocrinol 353(1-2): 109–113.

9. Csordás, G. r., et al. (2006). “Structural and functional features and significance of the physical linkage between ER and mitochondria.” Journal of Cell Biology 174(7): 915–921.

10. Cui, J., et al. (2004). “Regulation of the type 1 inositol 1,4,5-trisphosphate receptor by phosphorylation at tyrosine 353.” J Biol Chem 279(16): 16311–16316.

11. Eisner, V., et al. (2018). “Mitochondrial dynamics in adaptive and maladaptive cellular stress responses.” Nature Cell Biology 20(7): 755–765.

12. Filadi, R., et al. (2017). “The endoplasmic reticulum-mitochondria coupling in health and disease: Molecules, functions and significance.” Cell Calcium 62: 1–15.

13. Gabriel, B. M., et al. (2021). “Disrupted circadian oscillations in type 2 diabetes are linked to altered rhythmic mitochondrial metabolism in skeletal muscle.” Sci Adv 7(43): eabi9654.

14. Giorgi, C., et al. (2015). “Mitochondria-associated membranes: composition, molecular mechanisms, and physiopathological implications.” Antioxid Redox Signal 22(12): 995–1019.

15. Giorgi, C., et al. (2018). “The machineries, regulation and cellular functions of mitochondrial calcium.” Nature Reviews Molecular Cell Biology 19(11): 713–730.

16. Horner, S. M., et al. (2015). “Proteomic analysis of mitochondrial-associated ER membranes (MAM) during RNA virus infection reveals dynamic changes in protein and organelle trafficking.” PLoS One 10(3): e0117963.

17. Jagannathan, M., et al. (2010). “Toll-like receptors regulate B cell cytokine production in patients with diabetes.” Diabetologia 53: 1461–1471.

18. Karihtala, P. and Y. Soini (2007). “Reactive oxygen species and antioxidant mechanisms in human tissues and their relation to malignancies.” Apmis 115(2): 81–103.

19. Kim, J. A., et al. (2008). “Role of mitochondrial dysfunction in insulin resistance.” Circ Res 102(4): 401–414.

20. Lackner, L. L. (2019). “The Expanding and Unexpected Functions of Mitochondria Contact Sites.” Trends Cell Biol 29(7): 580–590.

21. Ma, J. H., et al. (2017). “Comparative Proteomic Analysis of the Mitochondria-associated ER Membrane (MAM) in a Long-term Type 2 Diabetic Rodent Model.” Scientific reports 7(1): 2062–2062.

22. Marchi, S., et al. (2017). “Endoplasmic Reticulum-Mitochondria Communication Through Ca(2+) Signaling: The Importance of Mitochondria-Associated Membranes (MAMs).” Adv Exp Med Biol 997: 49–67.

23. Missiroli, S., et al. (2018). “Mitochondria-associated membranes (MAMs) and inflammation.” Cell Death & Disease 9(3): 329.

24. Missiroli, S., et al. (2020). “The Role of Mitochondria in Inflammation: From Cancer to Neurodegenerative Disorders.” Journal of clinical medicine 9(3): 740.

25. Mohanty, A., et al. (2019). “Mitochondria: the indispensable players in innate immunity and guardians of the inflammatory response.” Journal of cell communication and signaling 13(3): 303–318.

26. Muneer, M. (2021). “Hypoglycaemia.” Adv Exp Med Biol 1307: 43–69.

27. Petrenko, V., et al. (2020). “In pancreatic islets from type 2 diabetes patients, the dampened circadian oscillators lead to reduced insulin and glucagon exocytosis.” Proc Natl Acad Sci U S A 117(5): 2484–2495.

28. Phillips, M. J. and G. K. Voeltz (2016). “Structure and function of ER membrane contact sites with other organelles.” Nat Rev Mol Cell Biol 17(2): 69–82.

29. Pizzino, G., et al. (2017). “Oxidative Stress: Harms and Benefits for Human Health.” Oxid Med Cell Longev 2017: 8416763.

30. Resende, R., et al. (2020). “Mitochondria, endoplasmic reticulum and innate immune dysfunction in mood disorders: Do Mitochondria-Associated Membranes (MAMs) play a role?” Biochimica et Biophysica Acta (BBA) - Molecular Basis of Disease 1866(6): 165752.

31. Rieusset, J. (2018). “Mitochondria-associated membranes (MAMs): An emerging platform connecting energy and immune sensing to metabolic flexibility.” Biochemical and Biophysical Research Communications 500(1): 35–44.

32. Rieusset, J. (2018). “The role of endoplasmic reticulum-mitochondria contact sites in the control of glucose homeostasis: an update.” Cell Death & Disease 9(3): 388.

33. Schieber, M. and Navdeep S. Chandel (2014). “ROS Function in Redox Signaling and Oxidative Stress.” Current Biology 24(10): R453–R462.

34. Sun, Y. and S. Ding (2020). “ER-Mitochondria Contacts and Insulin Resistance Modulation through Exercise Intervention.” International journal of molecular sciences 21(24): 9587.

35. Tubbs, E., et al. (2018). “Disruption of Mitochondria-Associated Endoplasmic Reticulum Membrane (MAM) Integrity Contributes to Muscle Insulin Resistance in Mice and Humans.” Diabetes 67(4): 636–650.

36. Volpe, C. M. O., et al. (2018). “Cellular death, reactive oxygen species (ROS) and diabetic complications.” Cell Death Dis 9(2): 119.

37. Wei, J. P., et al. (2018). “Research Progress on Non-Drug Treatment for Blood Glucose Control of Type 2 Diabetes Mellitus.” Chin J Integr Med 24(10): 723–727.

38. Xu, L., et al. (2022). “Macrophage Polarization Mediated by Mitochondrial Dysfunction Induces Adipose Tissue Inflammation in Obesity.” International journal of molecular sciences 23(16).

39. Yang, M., et al. (2020). “Mitochondria-Associated ER Membranes - The Origin Site of Autophagy.” Front Cell Dev Biol 8: 595.

40. Yang, S., et al. (2020). “Mitochondria-Associated Endoplasmic Reticulum Membranes in the Pathogenesis of Type 2 Diabetes Mellitus.” Front Cell Dev Biol 8: 571554.

41. Zorov, D. B., et al. (2014). “Mitochondrial reactive oxygen species (ROS) and ROS-induced ROS release.” Physiol Rev 94(3): 909–950.

